# Degenerative expansion of a young supergene

**DOI:** 10.1101/326645

**Authors:** Eckart Stolle, Rodrigo Pracana, Philip Howard, Carolina I. Paris, Susan J. Brown, Claudia Castillo-Carrillo, Stephen J. Rossiter, Yannick Wurm

## Abstract

***Suppressed recombination ultimately leads to gene loss, as demonstrated by the depauperate Y chromosomes of long-established XY pairs. To understand the shorter term effects, we used high-resolution optical mapping and k-mer distribution analysis on a young non-recombining region of fire ant social chromosomes. Instead of shrinking, the region has increased in length by more than 30%. This demonstrates that degenerative expansion can occur during the early evolution of non-recombining regions.***

---

Recombination facilitates the removal of deleterious mutations and creates advantageous combinations of alleles. However, in some circumstances reduced recombination is favored. This occurs during the early evolution of supergenes, in which selection favors the suppression of recombination between haplotypes with advantageous combinations of alleles at different loci^1,2^. Sex chromosomes harbor the most well-known supergene regions, but we now know that such genetic architectures are not rare and control variation for many other complex phenotypes^3,4^. Reduced recombination leads to reduced efficacy of selection, including reduced ability to remove deleterious mutations, because of interference among linked loci. This phenomenon^5^ is strongest in supergene variants where recombination is fully suppressed. For example, because the Y (or W) chromosome does not occur in the homozygous state, genetic hitchhiking and background selection affect the entire length of its supergene region^2,5–8^. This results in the gradual degeneration of Y (and W) chromosomes, with two striking long-term effects: the loss of protein-coding genes and the accumulation of repetitive elements^97,9^, both reducing gene density. This is particularly visible in the human Y chromosome which has approximately 14 times fewer genes and 5 times lower gene density than the X chromosome^10,11^.

Accumulation of repeats can already happen at early stages of Y chromosome evolution as shown in *Drosophila miranda* (age 1 million years, i.e. 10 million generations) and *Silene latifolia* (age 6 million years, i.e. 6 million generations)^12^. Intriguingly, the supergene region of suppressed recombination on the hermaphrodite determining Y^h^ chromosome of papaya (7 million years old, i.e. 7 million generations) is approximately two-fold larger than the homologous region in the X chromosome^13^. Such results suggests that large-scale accumulation of repetitive elements could precede gene loss. Because repetitive regions are difficult to study, we know little about how and when such “degenerative expansion”^14^ occurs.

The young social chromosome supergene system of the red fire ant *Solenopsis invicta* provides the opportunity to examine the early effects of restricted recombination. Two social chromosome supergene variants, SB and Sb, control a complex social phenotypic dimorphism where colonies have either one or up to dozens of reproductive queens^15,16^. The accumulation of unique SNP alleles indicates that recombination between the two variants has been suppressed for >350,000 years (i.e., >175,000 generations) over a chromosomal region encompassing >20 Mb and containing >400 protein-coding genes^16^. The suppression of recombination in *Bb* individuals has led to differentiation between SB and Sb throughout the entire length of the region^17^. SB can recombine in homozygote diploid *BB* queens. However, *bb* queens are never observed, either because they fail to reproduce, or because they die due to other intrinsic reasons^18^. Because Sb has no opportunity to recombine it should be affected by reduced efficacy of selection in a similar way to a Y or W chromosome.

To test whether degenerative expansion is an early effect of suppressed recombination, we apply a dual approach based on Bionano Genomics Irys optical mapping and short read Illumina sequence data. In a first step we optically mapped one haploid fire ant male carrying the SB variant and one carrying the Sb variant. For each individual, we created a *de novo* assembly of optical contigs (Suppl. File 1).

We first performed pairwise alignments between the optical assemblies of the two individuals to identify large (≥3kb) insertions and deletions (indels). The 187 deletions in the *b* individual were homogeneously distributed among the 16 chromosomes according to chromosome size (X^2^_d.f.=15_ = 24.02, p = 0.07). However, the social chromosome which carries the supergene region was significantly enriched in insertions (Fig. 1a and b): this chromosome harbors 33.7% (55) of the 163 mapped insertions despite representing only 8.4% (29.61 Mb) of the superscaffolded genome (350.94 Mb; X^2^_d.f.=15_ = 152, p < 10^−23^). Similarly, the cumulative length of insertions on the social chromosome was 58.5% (1.43 Mb) of the cumulative length of all insertions (2.44 Mb), higher than would be expected if the insertions were homogeneously distributed across chromosomes. We then identified “overhangs”, unaligned regions that flank alignments between the optical assemblies of the *B* and the *b* individuals. Such overhangs either represent indels, highly divergent sequences, or are regions where an optical assembly is too fragmented. The cumulative amount of overhanging sequence indicates that the supergene region is 5.27 Mb larger in the *b* individual than in the *B* individual. This is a significantly greater difference than for chromosomes 1 to 15 (−1.43 to −0. 25 Mb, X^2^_d.f.=15_ = 83.25, Bonferroni-corrected p < 10^−14^). Combining the indels detected with both methods, the *b* variant of the supergene region is 31.27% longer (total length 26.37 Mb) than the *B* variant (20.9 Mb). This difference in length is likely to be an underestimate because we cannot detect indels in parts of the genome that are fragmented in both assemblies.

To corroborate our results, we obtained short read Illumina sequence data for one *B* and one *b* sample from each of five different *S. invicta* populations. We independently estimated genome size and the proportion of repetitive sequence in the genome of each sample using the distribution of 21-nucleotide k-mer sequences ^19^ (Suppl. File 1). Estimated genome sizes for *b* samples were 3.59% larger (95% confidence interval: 2.02% to 5.16%) than those of *B* samples (paired one-sided t-test: p < 0.002; Fig. 1c). Using a previous estimate that the supergene region is 4.5% of the genome^17^, and assuming that the difference in genome size between the *b* and *B* samples is entirely due to the increase in size of Sb in the supergene region, these data indicate that the Sb variant of the supergene is 79.8% (44.9% to 114.7%) larger than the SB variant. Furthermore, the genomes of the *b* samples included 4.55 % more repetitive sequence (0.52% to 8.57%) than the *B* samples (paired one-sided t-test: p < 0.018, Suppl. File 1).

**Fig. 1.**
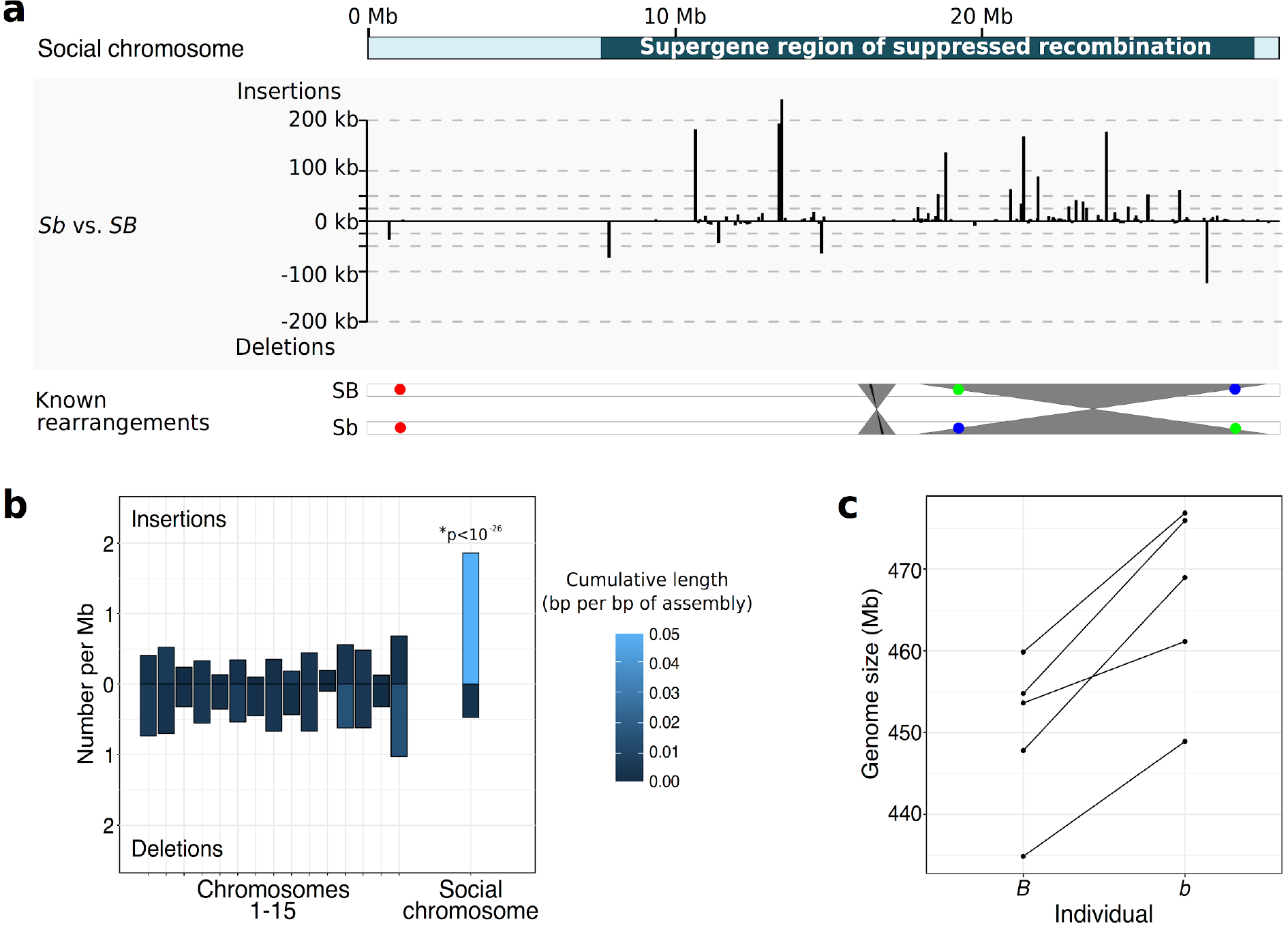
| Accumulation of insertions in *S. invicta* Sb supergene variant. **a**, Graph: Distribution of insertions and deletions along the social chromosome are largely within the supergene region (located from position 7.7 Mb to 28.6 Mb). Bottom: overview of known rearrangements between SB and Sb. Grey ribbons represent inversions detected in this study; black ribbon represents a previously known 48kb inversion; colored circles represent BAC-FISH markers A22, E17, E03^16^). **b**, Frequency and cumulative length of insertions and deletions in the pairwise comparison of optical contigs between an *S. invicta b* and an *S. invicta B* individual. Insertions were not homogeneously distributed among chromosomes (χ^2^_d.f.=15_ = 152, p < 10^−23^) with a significant enrichment exclusively on “social” chromosome 16, which carries the supergene region (Z-score = 11.1, Bonferroni-corrected p < 10^−26^). **c**, Genome sizes estimates using k-mer frequency distributions in raw Illumina sequence are higher in five *S. invicta b* individuals than in five paired *B* individuals.

Several close relatives of *S. invicta* are also socially polymorphic. Social polymorphism in *S. quinquecuspis* and *S. richteri* (common ancestry with *S. invicta* approximately 788,000 years ago; Suppl. File 1) is associated with the *Gp-9* locus which marks the supergene in *S. invicta*^20^. We created optical assemblies for one *Gp-9 B* sample and one *Gp-9 b* sample from each of the two relatives. To test whether *S. quinquecuspis* and *S. richteri* also carry the social supergene, we identified indels between the *B* and *b* optical assemblies in each of the three *Solenopsis* species. According to neighbor-joining trees based on presence and absence of indels for each of chromosomes 1 to 15, individuals cluster by species. In contrast, in a tree built using the region of the social chromosome supergene as known from *S. invicta*, the *b* individuals cluster separately from the *B* individuals. These data demonstrate that the supergene region exists in all three species, and that it likely has a single origin. These conclusions are further corroborated by inversions shared across species (Suppl. Files). The optical assemblies of the related species had lower contiguity than for *S. invicta* but provided the power to compare distributions of insertions and deletions. In both species, the supergene region in the *b* sample had a highly significant enrichment of insertions but not deletions in comparison to the *B* sample and to the rest of the genome (Suppl. File 1).

In sum, Sb has accumulated at least 30% more DNA content than SB. Despite extensive efforts^16,21^, we know of almost no differences in content of protein coding genes between Sb and SB. Our results thus show that Sb is in a stage of degenerative expansion in three *Solenopsis* species. This expansion is consistent with reduced efficacy of selection against mildly deleterious mutations such as insertions in non-coding regions^6,14,22^. Our study is the first to demonstrate degenerative expansion in an animal, but this may be a general feature of young nonrecombining regions. In stickleback fish, a nascent Y chromosome that is cytologically indistinguishable from the X includes Y-specific insertions and duplications^23^. Similarly, the older *Drosophila hydei* Y chromosome is smaller than its X chromosome counterpart, but carries some of the largest introns of the genome (≥3.6 Mb)^24^, perhaps a remnant of past chromosome-wide expansion. What type of selective pressure could be responsible for the absence of large deletions in a young supergene region such as the Sb supergene variant? Part of the answer could be that the system is too young, lacking mechanisms such as dosage compensation^2^ or gene relocation that could reduce the fitness cost of large deletions^8,14^. On the other hand, purifying selection could be much stronger in ants, where mutations are exposed to strong purifying selection in haploid males, than in organisms that are always diploid. Such strong purifying selection against loss of genetic material has indeed been observed in other organisms with haploid life stages^25,26^. Our study presents a striking example of degenerative expansion, providing evidence for this process as a hallmark of early supergene evolution and highlighting the important role of recombination (or the lack thereof) in shaping the genome.

## Methods

**Optical mapping.** For each of the of species *S. invicta*, *S. quinquecuspis* and *S. richteri*, we extracted high molecular weight (HMW) DNA from one haploid male pupae carrying the *B* genotype at the *Gp-9* locus and one carrying the *b* genotype, following the Bionano Genomics (BNG) IrysPrep animal tissue protocol (Suppl. files). Each sample was optically mapped using BNG nanochannel arrays for 30 cycles. Raw BNG Irys optical molecules ≥100 kb were processed, analysed and *de novo* assembled in IrysView (BNG, v2.4, scripts v5134, tools v5122AVX; Suppl. File 1).

**Optical assembly comparisons, optical chromosomes.** Comparisons between optical assemblies were performed by pairwise alignments using BNG IrysView (v2.4; Supplementary Methods). Large (≥3kb) insertions and deletions (indels) were detected as previously described^27^. A reciprocal alignment between *S. invicta* optical assemblies (b and B) yielded nearly identical results (95% of indel sites were recovered; data not shown), indicating high consistency of indel detection. We placed and oriented the optical contigs of the *S. invicta B* optical assembly onto the 16 linkage groups in the *S. invicta* genetic map^17^ using the alignment between the optical contigs and the scaffolds of the *S. invicta B* reference genome assembly^28^ (GCF_000188075.1; Suppl. File 1). The small portion of ambiguous placements of the optical contigs from this individual were resolved using information from optical contigs of the additional males.

**K-mer analyses of genome size, repeat content.** We sequenced five *B* and five *b S. invicta* haploid male individuals (*i.e.*, 5 pairs) on Illumina HiSeq4000 with 150 bp paired-end reads. After quality filtering and adapter trimming, we mapped cleaned reads to mitochondrion, phiX phage and *Wolbachia* reference genomes to enrich for *S. invicta* reads. From normalized numbers of the putative *S. invicta* reads, we used k-mer distribution analysis^19^ to estimate genome size and repeat content for each sample. Additionally, we subsampled 0.3× genome coverage of reads from each sample to characterise the types of repeats present^29^ (see Suppl. File 1).

**Phylogenetic analysis.** For phylogenetic tree reconstruction and dating we used mitochondrial sequences generated from Illumina short read data (Suppl. File 1). We additionally inferred phylogenetic relationships between samples based on presence and absence of shared indels detected in pairwise comparisons of at least 2 individuals in regions that had information (coverage) in all 6 individuals.

**Data availability.** The data sets generated and analysed during the current study are available as Suppl. File, NCBI (BioProject PRJNA397545: SUPPF_0000001241 - SUPPF_0000001246) and Genbank (accessions MF592128 - MF592133).

**Computer code.** Further details for specific analyses and software input files can be found in the Suppl. File 1. The source code of used scripts is available from github: https://github.com/estolle/BioNano-Irys-tools.

## Author contributions

E.S. and Y.W. conceived and designed the study. E.S., C.A.C.C. and C.I.P. sampled and identified and genotyped fire ants. E.S. and R.P. analysed genetic map, assembly inconsistencies and statistics. R.P. analysed dS ratios. E.S. P.H. and S.B. performed optical mapping, sample and data processing. E.S. did PCR-product and whole genome library preparation and sequencing, analysed phylogenies, repeats, sequence and optical mapping data. Y.W., E.S., R.P. and S.R. wrote the manuscript. All authors gave final approval for the publication.

## Acknowledgements

We thank Maria Cristina Arias, Susy Coelho, Diego Pereira Nogueira Da Silva, Nefertitis Curi, Natália Souza Araujo (Universidade de São Paulo, Brazil), Rodolfo Jaffé (Instituto Tecnólogico Vale, Belém Brazil), Emiliano Boné (Universidad de Buenos Aires, Argentina), Yanina Guillij (Dirección de Gestión de Usos Sustentables de los Recursos Naturales, Área Fauna y Flora Silvestre, Entre Ríos, Argentina), Dirección de Fauna Silvestre y Dirección de Ordenamiento Ambiental y Conservación de la Biodiversidad of Secretaria de Medio Ambiente of Argentina, Nazrath Nawaz, Thomas J. Colgan, Christoph Durrant, Christophe Eizaguirre, Mario dos Reis, Richard Nichols (Queen Mary University of London, UK), John Wang (Biodiversity Research Center, Academia Sinica, Taiwan), Michelle Coleman (Kansas State University, USA) and Bionano Genomics support staff for their help in organising and helping with sampling, permits, preparation, sequencing or analysis, useful discussions and comments on the manuscript.

This project was supported by the Deutscher Akademischer Austauschdienst (DAAD) Postdoc-Programm (E.S.: 570704 83) and European Commission Marie Curie Actions (E.S.: PIEF-GA-2013-623713). Additional support came from Biotechnology and Biological Sciences Research Council (Y.W. BB/K004204/1); the Natural Environment Research Council (grants NE/L00626X/1 (Y.W.), Strategic Capital Investment (S.R.)). Computing was performed at QMUL Research-IT, using NERC EOS Cloud and MidPlus computational facilities (QMUL VP Research fund and Engineering and Physical Sciences Research Council grant EP/K000128/1).

## Additional information

Supplementary Information 1: PDF containing additional figures and data.

